# Detection and genetic analysis of infectious spleen and kidney necrosis virus (ISKNV) in ornamental fish from non-clinical cases: First report from India

**DOI:** 10.1101/2020.08.12.247650

**Authors:** Sabyasachi Pattanayak, Anirban Paul, Pramoda Kumar Sahoo

**Affiliations:** National Referral Laboratory for Freshwater Fish Diseases, Fish Health Management Division, ICAR-Central Institute of Freshwater Aquaculture, Kausalyaganga, Bhubaneswar 751002, India

**Keywords:** ISKNV, Major capsid protein, Ornamental fish, PCR, Phylogeny

## Abstract

Infectious spleen and kidney necrosis virus (ISKNV), a type species of the genus *Megalocytivirus*, poses a threat to ornamental fish trade as most cases show nonspecific symptoms, thus making timely diagnosis challenging. Apparently health molly (*Poecilia sphenops*) and angelfish (*Pterophyllum scalare*) collected from two distinct geographic localities of India were screened for four genera under *Iridoviridae*, *Megalocytivirus* i.e, ISKNV, turbot reddish body iridovirus (TRBIV) and red seabream iridovirus (RSIV); ranaviruses and Singapore grouper iridovirus; and Lymphocystivirus through molecular approach. In total seven numbers out of 35 samples (20%), ISKNV genome fragments were detected. A PCR assay using major capsid protein (MCP) gene was standardised to detect and differentiate infections within the *Megalocytivirus* genus, even without aid of sequencing. This forms the first report of ISKNV infection in ornamental fish from India. Moreover, the ISKNV infection was confirmed by PCR and sequence analysis of MCP and ATPase genes. The sequence of these genes showed that Indian isolate being 99-100% similar to the complete genome or reference strain of ISKNV. Phylogenetic reconstruction demonstrated the present strain belonging to ISKNV genotype I. Furthermore, structural stability of the MCP revealed this strain was more stable than ISKNV genotype II, RSIV and TRBIV at 25°C and pH 7.0. Thus, strong pan-India surveillance is recommended to reduce trade risk.

## 1. Introduction

Ornamental fish culture and trade is widely accepted as an industry showing enormous potential and aggressive growth worldwide. In India, its popularity has boomed in recent times owing to its economic benefits due to a large internal market and the substantial unexplored potential for exports. The major groups of farm-bred ornamental fish are goldfish, barbs, tetras, swordtails, mollies, gourami, guppies, angelfish, fighting fish and platyfish in India. The molly (*Poecilia latipinna*) and angelfish (*Pterophyllum scalare*) are two crucial ornamental fish species, equally attractive and vibrantly coloured species, both of which can be kept as pets in confined spaces like aquaria or garden pools (Kumari et al., 2017). The total export of ornamental fish from India stood at around 42 tonnes, grossly estimated to have a value of approximately 8.4 crores (US $1.2 million) in 2017-18 alone (Singh, 2019).

On the flip side, the global ornamental fish trade poses a potential source for the spread of many exotic pathogens, particularly viruses causing both morbidity and mortality in fishes (Oyamatsu et al., 1997; George et al., 2015; Pragyan et al., 2019; Sahoo et al., 2016, 2020a, 2020b). Similarly, Sahoo et al. (2020b) investigated the infectious diseases in freshwater aquaculture farms of eastern India and found that the incidences of viral infections are more in ornamental fish. To-date, viral infections viz., viral nervous necrosis (VNN) in freshwater aquarium fishes ‘goldfish and rainbow shark’ (Jithendran et al., 2011), Cyprinid herpes virus-2 (CyHV-2) in goldfish (Sahoo et al., 2016), carp edema virus (CEV) in koi carp (Swaminathan et al., 2016; Sahoo et al., 2020a), ranavirus in koi carp (George et al., 2015) and marine ornamental similar damselfish (Sivasankar et al., 2017) have been reported from Indian ornamental fishes.

The family *Iridoviridae* consists of a large group of enveloped viruses possessing double-stranded DNA (Jancovich, 2012), which are known to cause large scale mortalities in fish. Taxonomists have divided this family into five genera viz. *Megalocytivirus*, *Iridovirus, Ranavirus*, *Lymphocystivirus* and *Chloriridovirus* (Kurita and Nakajima, 2012). Viruses belonging to the *Iridoviridae* family are typically vulnerable to both wild as well as freshwater fish, including ornamental fish. Iridoviruses capable of infecting fish have been accommodated in three major groups or genera viz., *Ranavirus*, *Lymphocystivirus* and *Megalocytivirus* (Chinchar et al., 2009). Since their discovery in the early 1990s, the group of Megalocytiviruses, in particular, have caused significant morbidity as well as financial losses in marine and freshwater fish culture, predominantly in Asia. Megalocytiviruses have been broadly distributed into three clusters, i.e., RSIV, infectious spleen and kidney necrosis virus (ISKNV) and turbot reddish body iridovirus (TRBIV), based on their major capsid protein (MCP) and ATPase gene phylogenetic analysis (Kurita and Nakajima, 2012), and recently another strain, three spine stickleback iridovirus (TSIV) have also been described (Murwantoko et al., 2018). He et al. (1998) first isolated ISKNV, a species of the genus *Megalocytivirus*, from mandarin fish in China. Subsequently ISKNV has also been reported from Japan (Tanaka et al., 2014), Malaysia (Subramaniam et al., 2014), Australia (Mohr et al., 2015), USA (Subramaniam et al., 2016), Germany (Jung-Schroers et al., 2016), Vietnam (Dong et al., 2017), Africa (Ramires et al., 2019) and Thailand (Thanasaksiri et al., 2019) in both freshwater and marine ornamental fishes. Phylogenetic analysis of the *Megalocytivirus* MCP gene sequence revealed the existence of two genotypes I and II of ISKNV (Kurita and Nakajima, 2012; Dong et al., 2017; Murwantoko et al., 2018).

Ornamental fish viruses have inherent inability to exhibit growth in commonly used fish cell lines, thus making diagnosis difficult. ISKNV like isolates, mostly those from freshwater ornamental fish, are challenging to culture *in vitro* (Kurita and Nakajima 2012, Rimmer et al., 2016) and CEV also being reported not to grow in any studied cell lines (Swaminathan et al., 2016). On multiple occasions, viral infection may remain latent until adverse environmental conditions like poor quality of water, rough handling of fish, or overcrowding, activate the disease (Sahoo et al., 2016, 2020a). ISKNV poses a threat to ornamental fish trade because *Megalocytiviral* infections have non-specific symptoms, thus making timely diagnosis challenging (Jeong et al., 2006, 2008; Subramaniam et al., 2014; Zainathan et al., 2017) and persistence of sub-clinical detected level of virus at the retailer site (Rimmer et al., 2015). Viral diagnosis based on candidate gene PCR-sequencing seems to be more accurate, reliable and faster than traditional phenotypic methods. Despite different molecular techniques like PCR, nested PCR, real-time PCR (Xu et al., 2008), and LAMP assay (Suebsing et al., 2016) being used for detection of ISKNV, the specificity for ISKNV has usually been low (Razak et al., 2014; Bobby et al., 2018) thereby failing to reliably differentiate ISKNV from other subgroups of *Megalocytiviruses*, i.e. RSIV and TRBIV. Hence, the current investigation was aimed at detecting the presence of *Megalocytiviruses* in two species of ornamental fishes, i.e., molly (*Poecilia sphenops*) and angelfish (*Pterophyllum scalare*) collected from two geographically distinct Indian ornamental fish markets from apparently healthy fish with a PCR based method even without aid of sequence confirmation. We standardised a precise and reliable nested PCR-based approach to detect ISKNV and segregate it from other species of *Megalocytiviruses*. Further this virus was also confirmed by molecular approach as described by OIE (OIE, 2019).

## 2. Materials and methods

### 2.1. Case history and sample collection

Thirty-five numbers of apparently healthy fish (Molly, *Poecilia sphenops* and angelfish, *Pterophyllum scalare*) were collected from seven different ornamental retailer shops of Kurla, Maharashtra and Bhubaneswar, Odisha states of India during the year 2018-2019 for routine screening of viruses in the National Referral Laboratory of Freshwater Fish Diseases at ICAR-Central Institute of Freshwater Aquaculture, Bhubaneswar, India. The fish were euthanized with overdose of MS222 (Sigma, USA), and liver, kidney, gills, brain and eye samples of individual fish were collected in 100% ethanol. The experiments were performed following the approval of the Institute Animal Ethics Committee.

### 2.2. Molecular screening for Megalocytiviruses

#### 2.2.1. DNA isolation

Tissue samples (~ 10 mg) were treated with proteinase K in lysis buffer (50 mM Tris/HCl, 100 mM NaCl, 100 mM EDTA, 1% [w/v] SDS, pH 8.0) and subjected to extraction with phenol/chloroform/isoamyl alcohol, followed by ethanol precipitation. The isolated DNA was diluted in TE (50 mM Tris/HCl, one mM EDTA, pH 7.5) buffer. Concentration and purity of the extracted DNA was determined by measuring OD at 260 and 280 nm using a NanoDrop ND1000 spectrophotometer (NanoDrop Technologies Inc., USA). The samples were stored at −20°C for further analysis. Before processing further, an equal amount of DNA samples from each organ of an individual fish were pooled to form one sample.

#### 2.2.2. PCR amplification

PCR was performed using five sets of published oligonucleotide primers for confirmation of ISKNV (two nested PCRs based on MCP gene and one PCR based on ATPase gene). Samples were also PCR screened for RSIV, TRBIV, EHNV, SGIV and LCDV using published primers and conditions (Table 1).

**Table 1.**
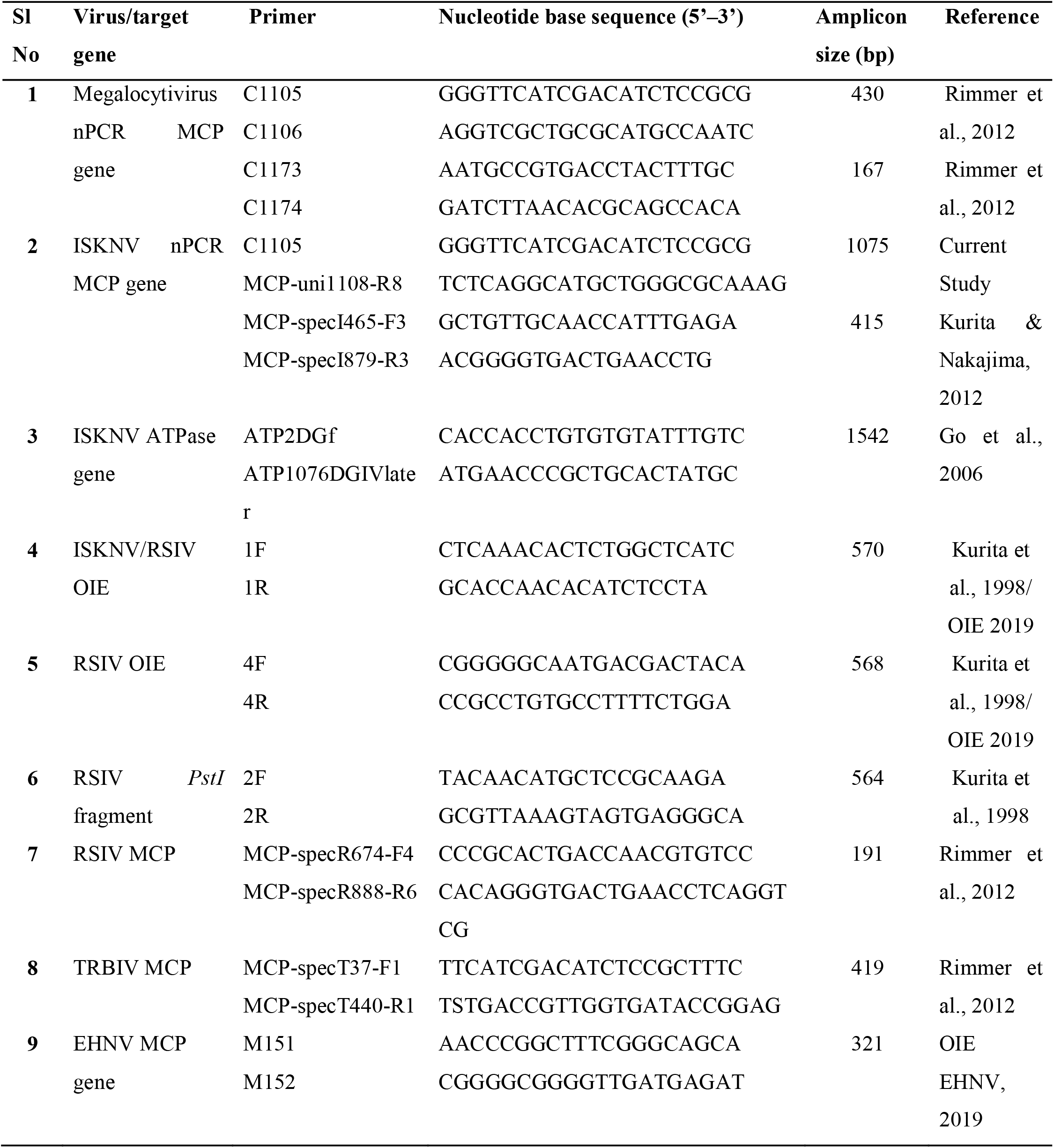
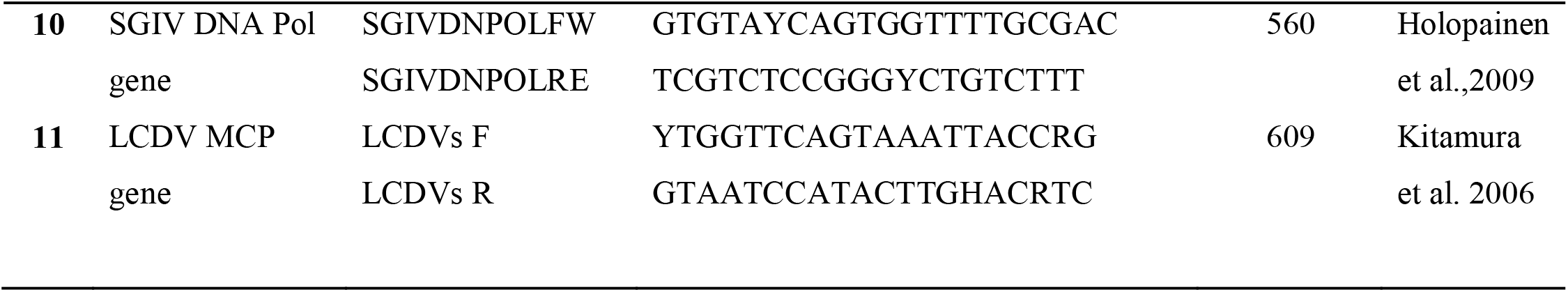
Details of the primers used in this study with expected size of amplicons

The samples were initially screened for MCP gene of MCV using two sets of published primers (Rimmer et al., 2012). In the first set, C1105 and C1106 (Table 1) primers were used in a final volume of 25 μL containing 1 μL of total DNA, 1.5 μL (10 pmol) of each primer, 0.25 μL of *Taq* DNA polymerase (5 U/μL), 2.5 μL of 10X *Taq* buffer A, 0.5 μL of dNTPs (2 mM) and ddH_2_O to make final volume to 25 μL. The reaction mix was subjected to 30 temperature cycles (30s at 94 °C, 30s at 55 °C and 1 min at 72 °C) after an initial denaturing step (15 min at 95 °C) followed by a final extension step of 7 min at 72 °C in a Veriti thermal cycler (Applied Biosystem). A nested PCR was performed using a second set primers C1073 and C1074 using similar conditions. These primers only confirm the presence of MCV at genus level.

After confirming the presence of MCV, samples were processed for molecular screening of ISKNV using a combination of published primers (Kurita and Nakajima, 2012; Rimmer et al., 2012) with partial modification. In the first step, primer C1105F and MCP-uni1108-R8 (unpublished pairing) were used in a final volume of 25 μL containing 1 μL of total DNA, 1.5 μL (10 pmol) of each primer, 0.25 μL of *Taq* DNA polymerase (5 U/μL1), 2.5 μL of 10X *Taq* buffer A, 0.5 μL of dNTPs (2 mM) and ddH_2_O to make final volume to 25 μL. The reaction mix was subjected to 35 temperature cycles (60 sec at 95 °C, 60 sec at 57 °C and 60 sec at 72 °C) after an initial denaturing step (300 sec at 95 °C) followed by a final extension step of 300 sec at 72 °C in a Veriti thermal cycler to amplify genus MCV. Further, nested PCR was performed using ISKNV specific primers MCP-specI 465-F3 and MCP-specI 879-R3 using the above first step product to increase the sensitivity and specificity for detecting only ISKNV as primers don’t amplify RSIV and TRBIV (Kurita and Nakajima, 2012). The reaction mix was subjected to 35 temperature cycles (1 min at 95 °C, 1 min at 58 °C and 1 min at 72 °C) after an initial denaturing step (5 min at 95 °C) followed by a final extension step of 5 min at 72 °C.

The samples were also screened for ISKNV ATPase gene (Mohr et al., 2015). PCR was carried out in a final volume of 25 μl reaction mix, as mentioned above for the MCP gene. Amplification was programmed for a preliminary 15 min denaturation step at 95 °C, followed by 30 cycles of denaturation at 94 °C for 30 s, annealing at 55 °C for 30 s, extension at 72 °C for 90 s, and finally extension at 72 °C for 7 min (Mohr et al., 2015). Several target amplicons were generated to strengthen confirmation of ISKNV as it would be the first report from India.

2.3 As described by OIE (2019), the samples were screened for ISKNV *Pst-I* restriction fragment using primer 1F and 1R, followed by RSIV DNA polymerase gene using primers 4F and 4R (Kurita et al., 1998). PCR was carried out in a final volume of 25 μl reaction mix, as mentioned above for the MCP gene and PCR conditions were set as described by Kurita et al. (1998) and OIE (2019).

### Sequencing and phylogenetic study

The PCR amplicons (using primers C1105/C1106, C1073/C1074,MCP-specI465-F3/MCP-specI879-R3,; ATP2DGf/ATP1076DGIVlater and ISKNV/RSIV 1F/1R) from four positive samples were purified using QIAquick Gel Extraction kit, and purified products were commercially sequenced (AgriGenome Labs Pvt. Ltd, Kochi, India). The nucleotide sequences were analysed using the Basic Local Alignment Search Tool (BLAST) of NCBI (http://www.ncbi.nlm.nih.gov/blast) to find out the homology. The amino acid sequences of ISKNV MCP gene of different isolates were retrieved from the NCBI database and aligned with the amino acid sequences of Indian samples. Multiple alignments were performed with MEGA 6 by using the ClustalW algorithm (Tamura, 2013). Phylogenetic analysis of ISKNV MCP sequences was performed through Maximum Likelihood method available in MEGA 6, and the phylogenetic tree was constructed using the Maximum Likelihood method. The amino acid sequences of different Megalocytiviruses were aligned and differentiated with ISKNV. Meanwhile, the nucleic acid sequences of three subgroups of Megalocytivirus i.e., RSIV, ISKNV and TRBIV were aligned and compared.

### 2.4. Protein stability prediction

Along with the sequence study, I-MUTANT 2.0 server was used to predict the stability of the MCV major capsid protein sequences at pH 7.0 and temperature 25 °C (Capriotti et al., 2005). This software can evaluate the stability change upon single-site mutation, starting from the protein structure or the protein sequence. I-Mutant 2.0 correctly predicts whether the protein mutation stabilises or destabilises the protein when the protein sequence is available.

## 3. Results

### 3.4 Identification of the virus in samples

Among 35 samples screened, seven numbers (20%) of samples were found to be positive for ISKNV. For detection of *Megalocytivirus*, MCP gene was screened using primers C1105/C1106 and C1073/C1074, and amplicons of 431 bp and 168 bp were obtained in first and second step PCRs, respectively (Figs. 1a and 1b). Aimed at detection of ISKNV, the tissues were screened with primers C1105/MCP-uni1108-R8 and MCP-specI465-F3/MCP-specI879-R3 primers. The samples were screened for RSIV *Pst*I fragment gene were found to be negative (Fig 1c). In seven samples, amplified product of 415 bp of ISKNV MCP gene were obtained in the nested PCR (Fig. 2a). Further, in same samples ISKNV ATPase gene were also amplified with amplicon size of 1542 bp using ATP2DGf and ATP1076DGIVlater primers (Fig. 2b). Meanwhile, the same samples amplified a specific band of 570 bp for *Pst*I gene by 1F and 1R (Fig. 2c) and no amplification of RSIV specific DNA polymerase gene by 4F and 4R, which indicated the presence of ISKNV infection in the samples. All the samples collected for this study were found to be negative for RSIV, TRBIV, EHNV, SGIV and LCDV using respective primer sets.

**Fig: 1a.**
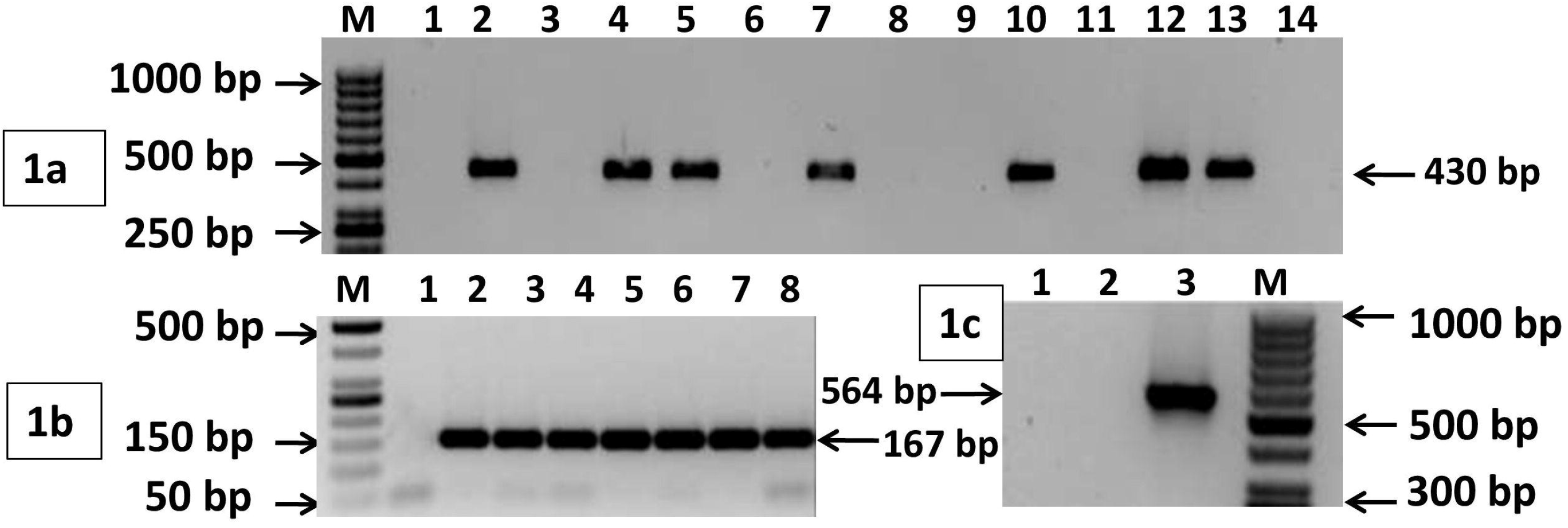
Samples amplified with C1105 and C1106 primers, with expected product sizes of 431 bp. Lane M represents 50 bp Ladder (Thermo Scientific); Lane 2 represent negative control; Lanes 3-15 represents fish samples, **1b.** Samples amplified with C1173 and C1174 primers; with expected product sizes of 167 bp. Lanes M represents 50 bp Ladder (Thermo Scientific) Lanes 2 represent negative control; Lanes 3-9 represents fish samples, **1c.** Samples did not amplified with RSIV 2F and 2R primer at expected size 564bp; Lane 1 represent negative control; Lane 2 represent fish sample; Lane 3 represent RSIV Positive control (RSIV Kag YT-96 DNA received from Prof Y. Kawato, NRIA, JFREA, Mie, Japan); Lane M represent 50 bp Ladder (Thermo Scientific)

**Fig: 2a.**
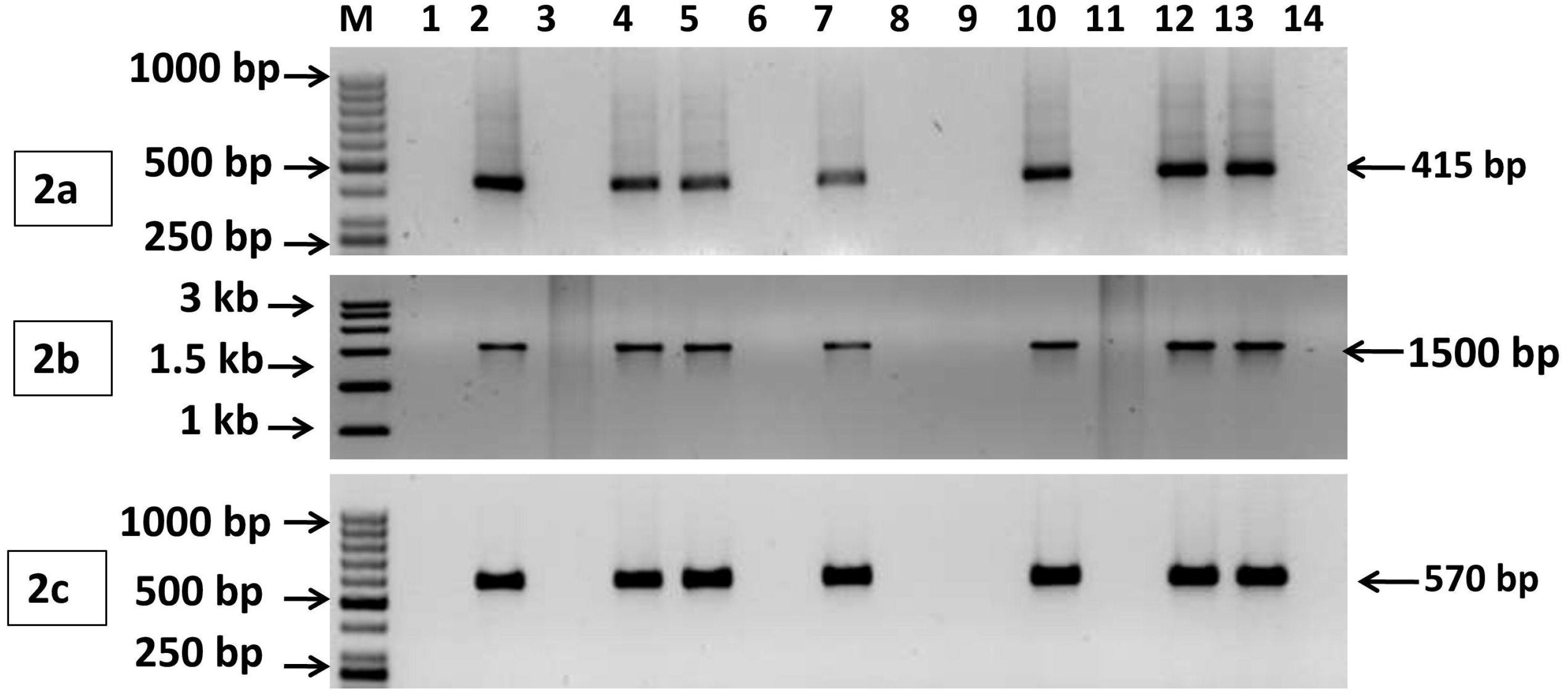
Samples amplified with MCP-specI465-F3 and MCP-specI879-R3 primers, with expected product sizes of 415 bp, Lane M represent 50 bp Ladder (Thermo Scientific), Lane 2 represents negative control and Lanes 3-15 represents fish samples; **2b.** Samples amplified with ATP2DGf and ATP1076DGIVlater primers; with expected product sizes of 1542 bp. Lane M represent 50 bp Ladder (Thermo Scientific), Lane 2 represents negative control and Lanes 3-15 represents fish samples; **2c.** Samples amplified with ISKNV/RSIV OIE 1F and 1R primer at expected size 570bp, Lane M represent 50 bp Ladder (Thermo Scientific), Lane 2 represent negative control and Lanes 3-15 represents fish samples.

### 3.5. Sequencing, comparison of MCP gene and phylogenetic study

ISKNV MCP gene amplicons obtained (using primers pairs C1105/C1106 and MCP-specI465-F3/MCP-specI879-R3) were sequenced, and a BLAST search of the sequence revealed 99% and 100% similarities with known previously published MCP genes of *Megalocytivirus* and ISKNV, respectively. Megalocytivirus MCP gene (primer C1105/C1106) sequence amplicon of [431 bp encoding 143 amino acids has been submitted to GenBank (GenBank accession number: MK084827). The 431 bp MCP gene fragment was found to be 99.77% similar with the complete genome or reference strain of ISKNV (AF371960) along with other ISKNV isolates (MK757444, KX354220, KT781098, AB666344, AB666337, AF370008), as well as ISKNV like viruses i.e., giant seaperch iridovirus isolate GSIV (JF264350) and African lampeye iridovirus (AY285745, AB109368) (Kurita and Nakajima, 2012). Also, the obtained sequence is 95.59% similar to RSIV (AB666327) and TRBIV (HM596017). Sequence alignment revealed a silent mutation observed at 399 bp position (GGCGly to GGTGly) in the MCP gene of Indian isolate (Fig. 3). Despite changes in the nucleotide sequence, the multiple 143 amino acid alignments of the MCP gene have shown 100% similarity with all the sequences mentioned (Fig. 4).

**Fig: 3.**
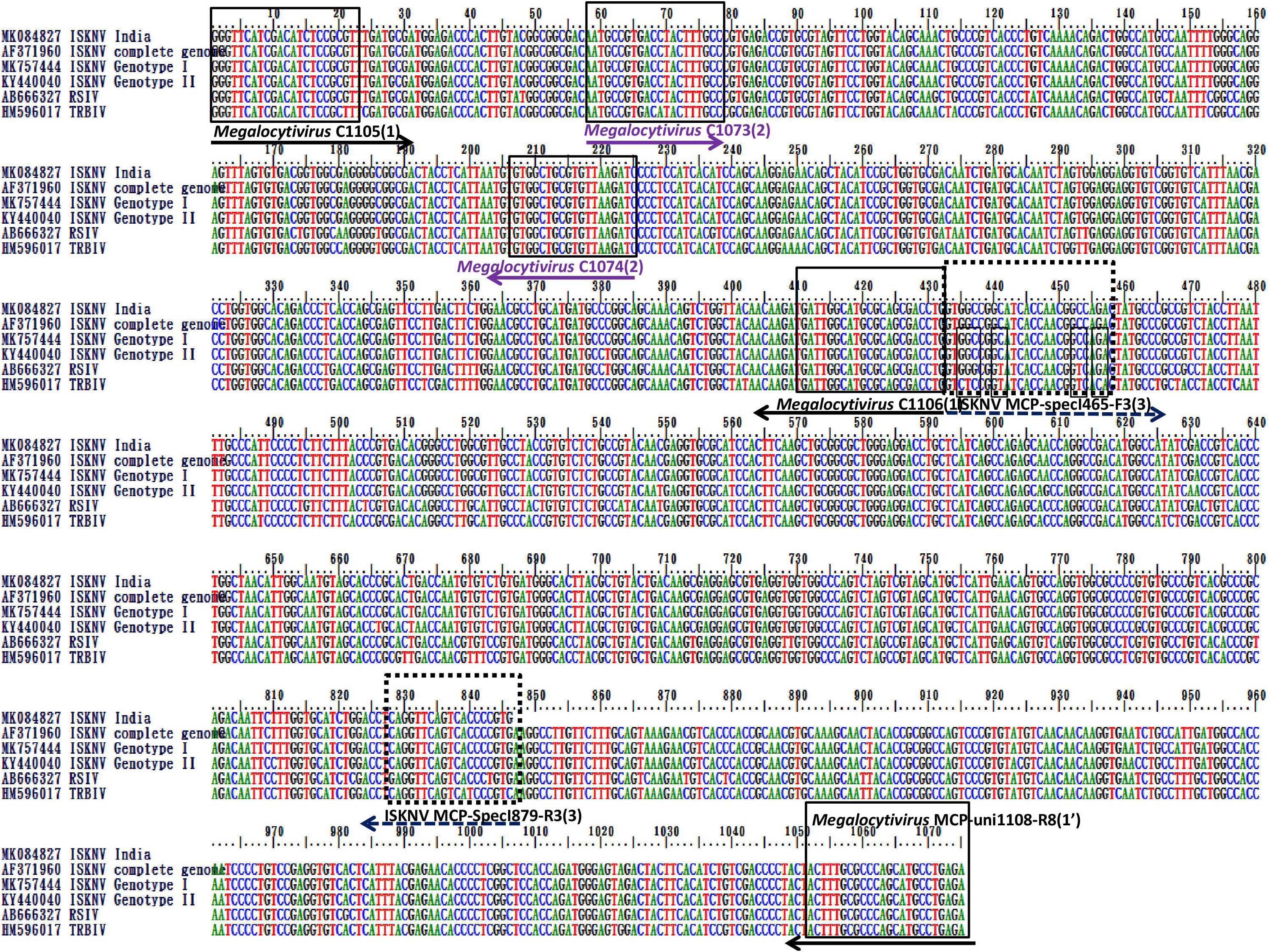
Nucleotide sequence alignments of primer sets C1105/C1106, C1073/C1074, C1105/MCP-uni1108-R8 and MCP-specI465-F3/MCP-specI879-R3 with major capsid protein (MCP) gene of infectious spleen and kidney necrosis virus (ISKNV) sequence of Indian isolate (accession: MK084827/MN518863), other published ISKNV isolates (accession: AF371960, MK757444, KY440040), red seabream iridovirus (RSIV) sequence (accession: AB666327) and turbot reddish body iridovirus (TRBIV) sequence (accession: HM596017). The nucleic acid variations are outlined in the ISKNV specific primer MCP-specI465-F3 alignment. The alignment result was obtained by graphic view of BioEdit Sequence Alignment Editor.

**Fig: 4.**
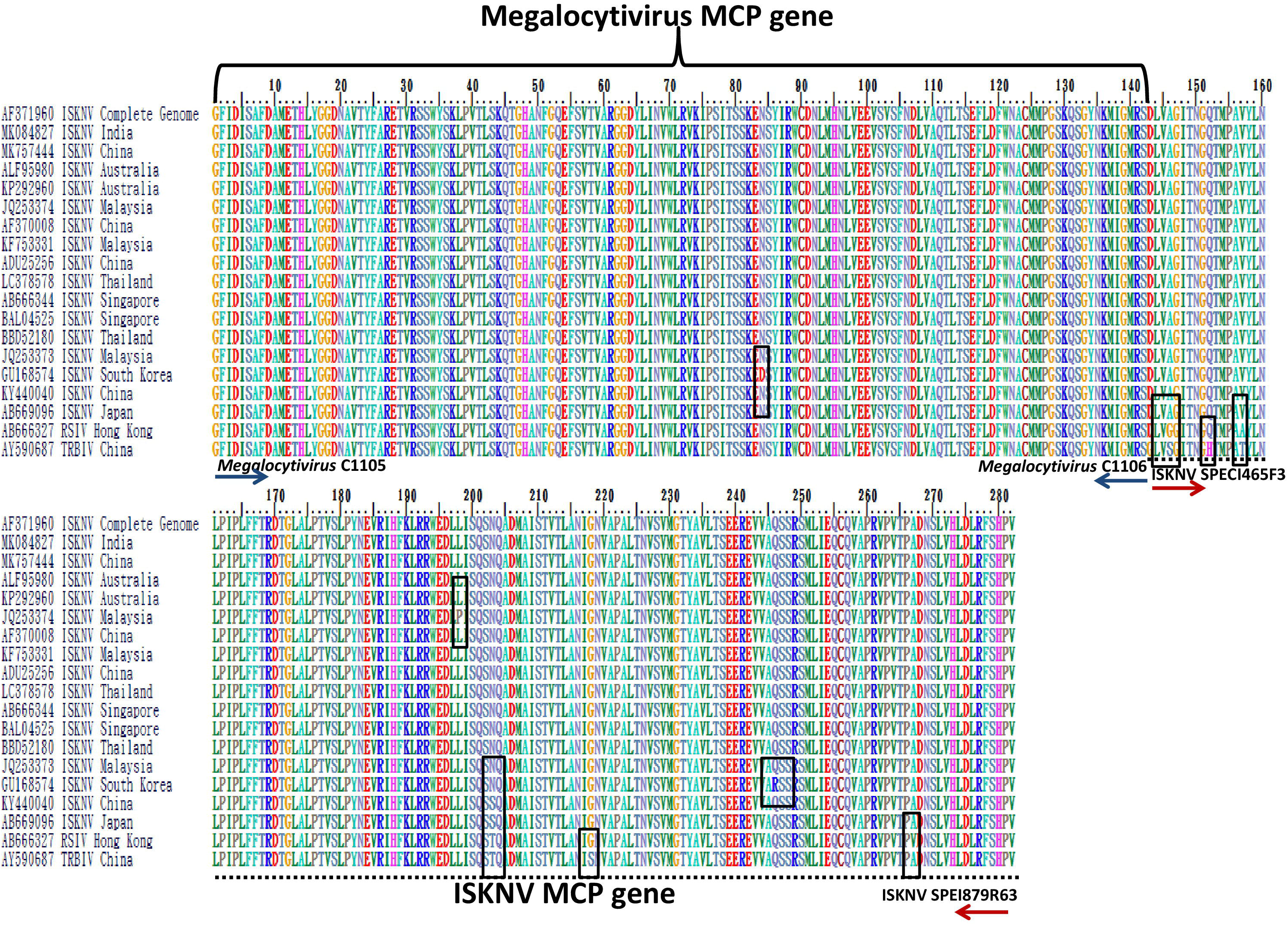
Amino acid alignment of MCP sequences (1-282) of different ISKNV isolates including India, RSIV and TRBIV isolates retrieve from NCBI. The amino acid variations are outlined. The alignment result is obtained by graphic view of Bio Edit Sequence Alignment Editor.

The amplicon of◻ 415 bp, encoding 138 amino acids of MCP gene sequence was found to be specific for ISKNV. It has been subsequently submitted to GenBank (GenBank accession number: MN518863). Both of these gene fragments (first 431 bp specific for *Megalocytivirus*, and next 415 bp specific for ISKNV) were combined to form a total of 846 nucleotides and 282 amino acids (Figs. 3, 4). The primer binding sites of *Megalocytivirus* specific primer C1105/C1106 and C1073/C1074 are common in the MCP gene of ISKNV, RSIV and TRBIV. To increase the sensitivity and confirmative diagnosis of ISKNV, the nested PCR using ISKNV specific MCP-specI465-F3 and MCP-specI879-R3 primer pair was successfully performed, that could differentiate ISKNV from the other *Megalocytivirus* species by amplifying only ISKNV (415 bp). Further the diagnosis was more validated based on *in silico* analysis for all three *Megalocytivirus* sequence primer binding regions in this study and also based on earlier report of Kurita and Nakajima (2012) and Mohr et al. (2015) (Fig. 3).

A phylogeny tree was constructed based on the MCP gene amino acid sequence (MK084827). The Indian isolate in the present study was found to be ISKNV genotype I. It showed 100% MCP gene sequence similarity with that of China (MK757444, AF370008, ADU25256), Australia (ALF95980, KP292960), Malaysia (JQ253373, KF753331), Singapore (AB666344, BAL04525), and Thailand (BBD52180, LC378578.1), and was clustered within the same clade in the phylogeny tree (Fig. 5). There were variations at the positions corresponding to amino acids 84 (Asn-Asp) and 198 (Leu-Pro) in the MCP sequences of South Korea (GU168574) and Malaysia (JQ253374), respectively (Fig. 4). Even so, the later two strains clustered with the Indian strain within the same clade of ISKNV genotype I (Fig. 5). Meanwhile, two other isolates of ISKNV, i.e., Japan (AB669096) and China (KY440040) differed from other ISKNV at amino acid position 203 (Asn-Ser) and were found to have clustered together as ISKNV genotype II (Figs. 4, 5). The two subspecies of Megalocytivirus, i.e., RSIV (AB666327) and TRBIV (AY590687) were segregated from ISKNV by amino acid position 146 (Ala-Gly), 157 (Val-Ala), 167 (Ala-Val) and 143 (Asp-Gly), 146 (Ala-Ser), 152 (Gln-His), 157 (Val-Thr), and 218 (Gla-Ser), respectively, and clustered at different clades (Figs. 4, 5).

**Fig: 5.**
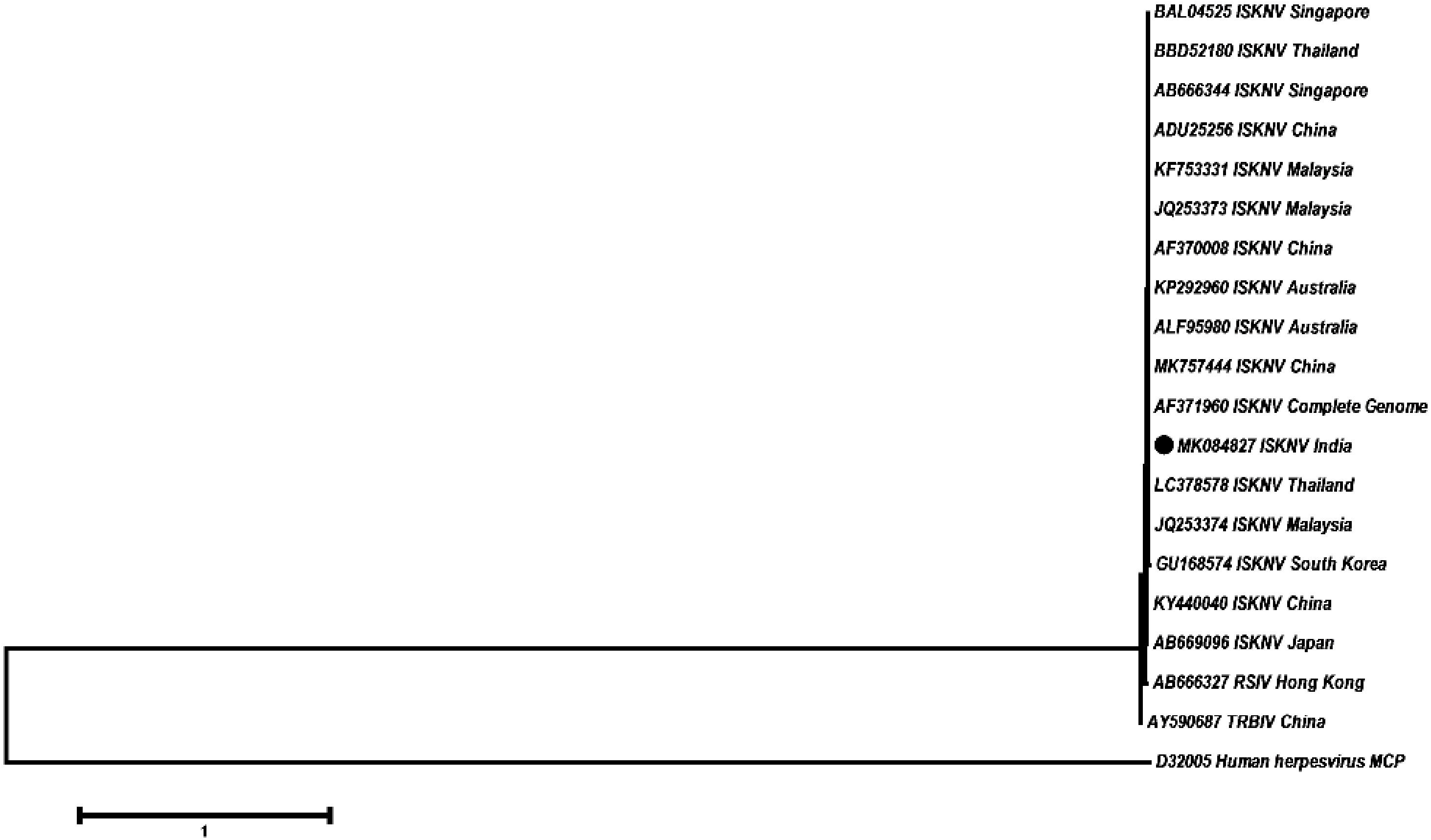
Phylogenetic tree based on the MCP gene amino acid sequence of ISKNV, RSIV and TRBIV. The tree was generated by MEGA 6 using Maximum Likelihood method of Clustal W 2. 1. MCP gene of human herpes virus was used as an outgroup.

For further confirmation, the ATPase and *Pst*I genes amplicons obtained using primers pairs (ATP2DGf/ATP1076DGIVlater and RSIV/ISKNV OIE 1F/1R) were sequenced, and a BLAST search of the sequence revealed 99% similarities with known previously published ATPase and laminin-like protein genes of ISKNV.

### 3.6. Prediction of protein stability of MCP

The stability of the MCP of ISKNV of Indian strain was compared *in silico* with ISKNV, RSIV and TRBIV isolates using I-Mutant software. The stability of MCP was found to reduce due to amino acid changes in ISKNV isolates of South Korea (84; N-D, 246; Q-R), Malaysia (198; L-P), Japan and China (Genotype II) (203; N-S) (Supplementary Fig. 1). MCP stability study indicated ISKNV MCP of having relatively better structural stability as compared to that of RSIV, whereas TRBIV MCP showed further decreased stability at 25°C and pH 7.0 (Supplementary Fig. 2).

## 4. Discussion

To our knowledge, it seems to be the first report of ISKNV infection in traded ornamental fish in India. The study further highlights a new combinational primer set to undertake a nested PCR (using C1105F/ MCP-uni1108-R8 and its nested MCP-specI465-F3 and MCP-specI879-R3) to directly identify ISKNV in PCR without ambiguity in one go, thereby delineating needs for further immediate need of sequence confirmation. Further, an increase in the sensitivity and specificity of only ISKNV-specific primer set published earlier (Kurita and Nakajima, 2012) was obtained to detect the virus in sub-clinical cases.. Moreover, the samples tested for RSIV and TRBIV, were found to be negative thus indicating ISKNV as the associated virus in the processed samples. The samples were reconfirmed by OIE based ISKNV detection protocol (Kurita et al., 1998, OIE, 2019).

The hosts affected by ISKNV are relatively wide-ranging; nevertheless, freshwater fish species are the predominantly affected species (Kurita and Nakajima, 2012; Sihu et al., 2017). Several outbreaks of ISKNV diseases in freshwater fishes have been reported in the ornamental fish of Germany (Jung-Schroers et al. 2016), Australia (Mohr et al. 2015; Rimmer et al. 2017), and Malaysia (Subramaniam et al. 2014; Zainathan et al. 2017) and more recently in Africa (Ramirez-Paredes et al., 2019). Detection of ISKNV in two ornamental fish species, molly (*P. sphenops*) and angelfish (*P. scalare*) obtained in this study were also reported earlier from other countries (Rodger et al., 1997; Rodger et al., 2003; Yanong and Waltzek, 2010; Go et al., 2016; Zainathan et al., 2017). Subramaniam et al. (2014) reported many positive cases of ISKNV in ornamental fishes which did not show any clinical signs. This further confirmed the fact that *Megalocytivirus* infects visibly healthy fish species; this condition could be termed persistent or asymptomatic infection as reported by Jeong et al. (2006, 2008). In comparison to clinically infected fish tissue, DNA concentration of *Megalocytivirus* was approximately 10^5^ times higher in asymptomatic fish, as proven by (Jeong et al., 2006). An earlier report finds the presence of ISKNV from apparently healthy fish at temperatures below 20 °C (He et al., 2002). When water temperature was 28 °C, juvenile Chinese perch challenged with ISKNV, displayed clinical signs with 100% mortality (He et al., 2000). It seems *Megalocytivirus* requires high temperature to multiply (Liu et al., 2019) that needs further study after isolation of virus. Further, recent report of emergence in causing mass mortality in tilapia in Africa raise a big concern for tilapia farming (Ramirez-Paredes et al., 2019), if not taken proper care, likely entry of it from ornamental fish to tilapia species would be disastrous.

Currently, the OIE Manual for Diagnostic Tests for Aquatic Animals does not identify a single test that would be suitable for accurate and specific detection of ISKNV (Kurita et al., 1998, OIE, 2018, 2019). Out of the recommended criteria for confirmatory diagnosis, conventional PCR detects the two listed genotypes, whereas others require specialized reagents (monoclonal antibodies) or are not applicable to all ISKNV like isolates (eg. virus isolation) (Johnson et al., 2019). Better methods remain to be identified for the specific diagnosis of ISKNV infections with particular reference to the differentiation of gene sequence, subclinical infections and clinical disease. Although there are several reports of screening the ISKNV virus by PCR and sequencing using a *Megalocytivirus* genus primer (C1105/C1106), none of these reports used ISKNV species specific primers described by Kurita and Nakajima (2012) and Mohr et al. (2015) (Go et al., 2006; Xu et al., 2008; Rimmer et al., 2012; Subramaniam et al., 2014; Razak et al., 2014; Mohr et al., 2015; Rimmer et al., 2017: Dong et al., 2017; Bobby et a., 2018; Johnson et al., 2019; Zainathan et al., 2019). Hence in this investigation, a nested PCR assay was standardised to detect and differentiate ISKNV infections within the *Megalocytivirus* genus of the family *Iridoviridae*.

The major capsid protein (MCP) gene is relatively conserved among the viruses of the family *Iridoviridae*, for which it can be considered as one of the most important genes for the analysis of the genetic relationships among this family (Williams et al., 1996; Tidona et al., 1998; Liu et al., 2019). The amino acid sequence analysis of *Megalocytivirus* specific MCP gene regions revealed 100% similarity with all the three *Megalocytivirus* subgroup. However, subgrouping issue was resolved by use of new combinational nested ISKNV PCR with internal primer set of ISKNV (Kurita and Nakajima, 2012), as it was 100% similar with only ISKNV genome with the produced amplicons, that emphasizes upon ISKNV subtyping with the current PCR system in one go.

ISKNV viruses can be further divided into two sub-clusters: genotypes I and II (Rimmer et al., 2012; Dong et al., 2017). Phylogenetic analysis revealed that the Indian isolate belonged to genotype I, which was more closely related to ISKNV isolates of China (MK757444, AF370008, ADU25256), Australia (ALF95980, KP292960), Malaysia (JQ253373, KF753331), Singapore (AB666344, BAL04525), and Thailand (BBD52180, LC378578.1), thus forming a single cluster. This isolate formed a different cluster with Japan (AB669096) and China (KY440040) isolates which belonged to ISKNV genotype II (Dong et al., 2017).

A proteomic study of ISKNV demonstrated that ISKNV virions possess 38 viral structural proteins, where MCP is the main structural component of the virus particle, comprising 40 to 45% of the total particle polypeptide (Dong et al., 2011; Jia et al., 2013). Jia et al. (2013) reported first time, ISKNV MCP interacted with Mandarin fish caveolin 1 (mCav-1), which gives an evidence of a viral structural protein interacted with a host protein in the family *Iridoviridae* for its primary infection. This information indicates that, major capsid proteins play an essential role in the ISKNV infection by virus-host protein interactions. Further study can help us to understand the mechanisms of viral pathogenicity, which may be related to the structural proteins and virus-induced immunological responses. Keeping in view of the above study, we investigated structural stability of the MCP of ISKNV, RSIV and TRBIV using I-Mutant software, and it was found that the ISKNV is more stable at 25°C and pH 7.0. Among different ISKNV isolates, Indian isolate along with similar MCP sequence isolates were found more structurally stable. Furthermore, ISKNV genotype I (Indian isolate) of MCP was found to be having more structural stability than ISKNV genotype II. This study suggests that ISKNV genotype I might be more virulent, but further investigation is needed to establish its probable role in influencing viral pathogenicity at different water temperatures or stressors.

To summarize, it seems to be the first report of ISKNV from ornamental fish meant for either internal or external trade. The current investigation also established an improved molecular diagnostic tool for detection of ISKNV pending sequencing confirmation, even from sub-clinical cases. Given the promiscuous nature of Megalocytiviruses, a strict biosecurity seems to be essential for its further spread to cultured fish species and also equally important to go for pan-India surveillance for this virus.

## Acknowledgements

The authors wish to thank the Director, ICAR-Central Institute of Freshwater Aquaculture, Kausalyaganga, Bhubaneswar, India for providing necessary facilities during this study. Funding support form Department of Biotechnology, New Delhi is duly acknowledged.

## Data availability statement

The data that supports the findings of this study are available in the supplementary material of this article (supplementary Figs. 1 and 2).

---

**Supplementary Fig. 1.** Protein stability prediction of different ISKNV strain including Indian isolate. I-MUTANT 2.0 server was used to predict the stability of the protein sequences at pH 7.0 and temperature 25°C.

**Supplementary Fig. 2.** Protein stability prediction and comparison of stability upon mutation at pH 7.0 and temperature 25°C, among Indian ISKNV isolate with RSIV and TRBIV using I-MUTANT 2.0 server.

## References

Bermúdez, R., Losada, A.P., de Azevedo, A.M., Guerra-Varela, J., Pérez-Fernández, D., Sánchez, L., Padrós, F., Nowak, B., Quiroga, M.I., 2018. First description of a natural infection with spleen and kidney necrosis virus in zebrafish. J. Fish Dis. 41, 1283–1294.

Bobby, G., Hong, T.K., Addis, S.N.K., Wahid, M.E.A., Sung, Y.Y., Zainathan, S.C., 2018. First detection of Megalocytivirus in oysters (*Crassostrea iredalei*) from Marudu Bay, Sabah, Malaysia. Aquacult. Aquarium Conserv. Legis. 11, 1537–1547.

Capriotti, E., Fariselli, P., Casadio, R., 2005. I-Mutant 2.0., Predicting stability changes upon mutation from the protein sequence or structure. Nucleic Acids Res. [Online], 33 (Web Server issue), W306–310.

Chinchar, V.G., Hyatt, A., Miyazaki, T., Williams, T., 2009. Family Iridoviridae: poor viral relations no longer. Curr. Top. Microbiol. Immunol. 328, 123–170.

Dong, C.F., Xiong, X.P., Shuang, F., Weng, S.P., Zhang, J., Zhang, Y., Luo, Y.W., He, J.G., 2011. Global landscape of structural proteins of infectious spleen and kidney necrosis virus. J. Virol. 85, 2869–2877.

Dong, H.T., Jitrakorn, S., Kayansamruaj, P., Pirarat, N., Rodkhum, C., Rattanarojpong, T., Senapin, S., Saksmerprome, V., 2017. Infectious spleen and kidney necrosis disease (ISKND) outbreaks in farmed barramundi (*Lates calcarifer*) in Vietnam. Fish Shellfish Immunol. 68, 65–73.

George, M.R., John, K.R., Mansoor, M.M., Saravanakumar, R., Sundar, P., Pradeep, V., 2015. Isolation and characterization of a ranavirus from koi, *Cyprinus carpio* L., experiencing mass mortalities in India. J. Fish Dis. 38, 389–403.

Go, J., Lancaster, M., Deece, K., Dhungyel, O., Whittington, R., 2006. The molecular epidemiology of iridovirus in Murray cod (*Maccullochella peelii peelii*) and dwarf gourami (*Colisa lalia*) from distant biogeographical regions suggests a link between trade in ornamental fish and emerging iridoviral diseases. Mol. Cell. Probes. 20, 212–222.

Go, J., Waltzek, T.B., Subramaniam, K., Yun, S.C., Groff, J.M., Anderson, I.G., Chong, R., Shirley, I., Schuh, J.C.L., Handlinger, J.H., Tweedie, A., 2016. Detection of infectious spleen and kidney necrosis virus (ISKNV) and turbot reddish body iridovirus (TRBIV) from archival ornamental fish samples. Dis. Aquat. Org. 122, 105–123.

He, J.G., Zeng, K., Weng, S.P., Chan, S.M., 2000. Systemic disease caused by an iridovirus-like agent in cultured mandarin fish *Siniperca chuatsi* (Basillewsky), in China. J. Fish Dis. 23, 219–222.

He, J.G., Zeng, K., Weng, S.P., Chan, S.M., 2002. Experimental transmission, pathogenicity and physical–chemical properties of infectious spleen and kidney necrosis virus (ISKNV). Aquaculture 204, 11–24.

Holopainen, R., Ohlemeyer, S., Schütze, H., Bergmann, S.M., Tapiovaara, H., 2009. Ranavirus phylogeny and differentiation based on major capsid protein, DNA polymerase and neurofilament triplet H1-like protein genes. Dis. Aquat. Org. 85, 81–91.

Iwamoto, R., Hasegawa, O., LaPatra, S., Yoshimizu, M., 2002. Isolation and characterization of the Japanese flounder (*Paralichthys olivaceus*) lymphocystis disease virus. J. Aquat. Anim. Health 14, 114–123.

Jancovich, J.K., 2012. Family iridoviridae. Virus taxonomy: ninth report of the international committee on taxonomy of viruses, 193–210.

Jeong, H.D., Lyu, J.H., Jeong, J.B., Kim, H.Y., Jun, L.J., Cho, H.J., Lee, J.W., 2006. Detection and distribution of iridoviruses in five freshwater ornamental fish species. J. Fish Dis. 19, 197–206.

Jeong, J.B., Kim, H.Y., Jun, L.J., Lyu, J.H., Park, N.G., Kim, J.K., Do Jeong,H., 2008. Outbreaks and risks of infectious spleen and kidney necrosis virus disease in freshwater ornamental fishes. Dis. Aquat. Org. 78, 209–215.

Jia, K.T., Wu, Y.Y., Liu, Z.Y., Mi, S., Zheng, Y.W., He, J., Weng, S.P., Li, S.C., He, J.G., Guo, C.J., 2013. Mandarin fish caveolin 1 interaction with major capsid protein of infectious spleen and kidney necrosis virus and its role in early stages of infection. J. Virol. 87, 3027–3038.

Jithendran, K.P., Shekhar, M.S., Kannappan, S., Azad, I.S., 2011. Nodavirus infection in freshwater ornamental fishes in India: diagnostic histopathology and nested RT-PCR. Asian Fish. Sci. 24, 12–19.

Johnson, S.J., Hick, P.M., Robinson, A.P., Rimmer, A.E., Tweedie, A., Becker, J.A., 2019. The impact of pooling samples on surveillance sensitivity for the megalocytivirus, Infectious spleen and kidney necrosis virus. Transbound. Emerg. Dis. 66, 2318–2328.

Jung-Schroers, V., Adamek, M., Wohlsein, P., Wolter, J., Wedekind, H., Steinhagen, D., 2016. First outbreak of an infection with infectious spleen and kidney necrosis virus (ISKNV) in ornamental fish in Germany. Dis. Aquat. Org. 119, 239–244.

Kumari, A., Kumar, S., Kumar, A., 2017. Study of life compatibility and growth of selected ornamental fishes under aquarium in Sanjay Gandhi Biological Park. Int. J. Curr. Microbiol. App. Sci. 6, 3166–3172.

Kurita, J., Nakajima, K., 2012. Megalocytiviruses. Viruses 4, 521–538.

Kurita, J., Nakajima, K., Hirono, I., Aoki, T., 1998. Polymerase chain reaction (PCR) amplification of DNA of red seabream iridovirus (RSIV). Fish Pathol. 33, 17–23.

Liu, L., Yu, L., Fu, X., Lin, Q., Liang, H., Niu, Y., Li, N., 2019. First report of megalocytivirus (*Iridoviridae*) in cultured bluegill sunfish, *Lepomis macrochirus*, in China. Microb. Pathog. 135, 103617.

Mohr, P.G., Moody, N.J., Williams, L.M., Hoad, J., Cummins, D.M., Davies, K.R., Crane, M.S., 2015. Molecular confirmation of infectious spleen and kidney necrosis virus (ISKNV) in farmed and imported ornamental fish in Australia. Dis. Aquat. Org. 116, 103–110.

Murwantoko, Sari, D.W.K., Handayani, C.R., Whittington, R.J., 2018. Genotype determination of megalocytivirus from Indonesian marine fishes. Biodiversitas 19, 1730–1736.

OIE (World Organisation for Animal Health) (2018). Aquatic Animal Health Code, 21st edn. OIE, Paris.

OIE. (2019). Red sea bream iridoviral disease. In Manual of diagnostic tests for aquatic animals. Retrieved from https://www.oie.int/fileadmin/Home/eng/Health_standards/aahm/current/chapitre_rsbid.pdf

Oyamatsu T., Matoyama, H., Yamamoto, K., Fukuda, H., 1997. A trial for detection of carp edema virus by using polymerase chain reaction. Aqua. Sci. 45, 247–251.

Pragyan, D., Bajpai, V., Suman, K., Mohanty, J., Sahoo, P.K., 2019. A review of current understanding on carp edema virus (CEV): A threatful entity in disguise. Int. J. Fish. Aquat. Stud. 7, 87–93.

Ramires, G., Paley, R.K., Hunt, W., Feist, S.W., Stone, D.M., Field, T., Hayden, D., Ziddah, P.A., Duodu, S., Wallis, T., Verner-Jeffreys, D., 2019. First detection of infectious spleen and kidney necrosis virus (ISKNV) associated with massive mortalities in farmed tilapia in Africa. bioRxiv 680538, (doi: https://doi.org/10.1101/680538).

Razak, A.A., Ransangan, J., Sade, A., 2014. First report of Megalocytivirus (Iridoviridae) in grouper culture in Sabah, Malaysia. Int. J. Curr. Micro. Ap. Sc. 3, 896–909.

Rimmer, A.E., Becker, J.A., Tweedie, A., Lintermans, M., Landos, M., Stephens, F., Whittington, R.J., 2015. Detection of dwarf gourami iridovirus (infectious spleen and kidney necrosis virus) in populations of ornamental fish prior to and after importation into Australia, with the first evidence of infection in domestically farmed Platy (*Xiphophorus maculatus*). Prev. Vet. Med. 122, 181–194.

Rimmer, A.E., Becker, J.A., Tweedie, A., Whittington, R.J., 2012. Development of a quantitative polymerase chain reaction (qPCR) assay for the detection of dwarf gourami iridovirus (DGIV) and other megalocytiviruses and comparison with the Office International des Epizooties (OIE) reference PCR protocol. Aquaculture 358, 155–163.

Rimmer, A.E., Whittington, R.J., Tweedie, A., Becker, J.A., 2016. Susceptibility of a number of Australian freshwater fishes to dwarf gourami iridovirus (infectious spleen and kidney necrosis virus). J. Fish Dis. 40, 293–310.

Rodger, H.D., Kobs, M., Macartney, A., Frerichs, G.N., 1997. Systemic iridovirus infection in freshwater angelfish, *Pterophyllum scalare* (Lichtenstein). J. Fish Dis. 20, 69–72.

Sahoo, P.K., Pattanayak, S., Paul, A., Sahoo, M.K., Rajesh Kumar, P., 2020a. Carp edema virus in ornamental fish farming in India: A potential threat to koi carps but not to co-cultured Indian major carp or goldfish. Indian J. Exp. Biol. 58, 254–262.

Sahoo, P.K., Paul, A., Sahoo., M.K., Pattanayak, S,, Rajesh Kumar, P., Das, B.K., 2020b. Incidences of infectious diseases in freshwater aquaculture farms of eastern India: a passive surveillance based study from 2014-2018. J. Aquac. Res. Dev. 11, 1–5.

Sahoo, P.K., Swaminathan, T.R., Abraham, T.J., Kumar, R., Pattanayak, S., Mohapatra, A., Rath, S.S., Patra, A., Adikesavalu, H., Sood, N., Pradhan, P.K., 2016. Detection of goldfish haematopoietic necrosis herpes virus (Cyprinid herpesvirus-2) with multi-drug resistant *Aeromonas hydrophila* infection in goldfish: First evidence of any viral disease outbreak in ornamental freshwater aquaculture farms in India. Acta Trop. 161, 8–17.

Shiu, J.Y., Hong, J.R., Ku, C.C., Wen, C.M., 2018. Complete genome sequence and phylogenetic analysis of megalocytivirus RSIV-Ku: A natural recombination infectious spleen and kidney necrosis virus. Arch. Virol. 163, 1037–1042.

Singh, G., 2019. Pretty lucrative – India’s surge in ornamental fish farming. The Fish Site, 5m.

Sivasankar, P., John, K.R., George, M.R., Mageshkumar, P., Manzoor, M.M., Jeyaseelan, M.P., 2017. Characterization of a virulent ranavirus isolated from marine ornamental fish in India. Virus Dis. 28, 373–382.

Subramaniam, K., Gotesman, M., Smith, C.E., Steckler, N.K., Kelley, K.L., Groff, J.M., Waltzek, T.B., 2016. Megalocytivirus infection in cultured Nile tilapia *Oreochromis niloticus*. Dis. Aquat. Org. 119, 253–258.

Subramaniam, K., Shariff, M., Omar, A.R., Hair-Bejo, M., Ong, B.L., 2014. Detection and molecular characterization of infectious spleen and kidney necrosis virus from major ornamental fish breeding states in Peninsular Malaysia. J. Fish Dis. 37, 609–618.

Suebsing, R., Pradeep, P.J., Jitrakorn, S., Sirithammajak, S., Kampeera, J., Turner, W.A., Saksmerprome, V., Withyachumnarnkul, B., Kiatpathomchai, W., 2016. Detection of natural infection of infectious spleen and kidney necrosis virus in farmed tilapia by hydroxynapthol blue-loop-mediated isothermal amplification assay. J. Appl. Microbiol. 121(1), 55–67.

Swaminathan, T.R., Kumar, R., Dharmaratnam, A., Basheer, V.S., Sood, N., Pradhan, P.K., Sanil, N.K., Vijayagopal, P., Jena, J.K., 2016. Emergence of carp edema virus in cultured ornamental koi carp, *Cyprinus carpio* koi, in India. J. Gen. Virol. 97, 3392–3399.

Tamura, K., Stecher, G., Peterson, D., Filipski, A., Kumar, S., 2013. MEGA6: molecular evolutionary genetics analysis version 6.0. Mol. Biol. Evo. 30, 2725.

Tanaka, N., Izawa, T., Kuwamura, M., Higashiguchi, N., Kezuka, C., Kurata, O., Wada, S., Yamate, J., 2014. The first case of infectious spleen and kidney necrosis virus (ISKNV) infection in aquarium-maintained mandarin fish, *Siniperca chuatsi* (Basilewsky), in Japan. J. Fish Dis. 37, 401–405.

Thanasaksiri, K., Takano, R., Fukuda, K., Chaweepack, T., Wongtavatchai, J., 2019. Identification of infectious spleen and kidney necrosis virus from farmed barramundi *Lates calcarifer* in Thailand and study of its pathogenicity. Aquaculture 500, 188–191.

Tidona, C.A., Schnitzler, P., Kehm, R., Darai, G., 1998. Is the major capsid protein of iridoviruses a suitable target for the study of viral evolution? Virus Gene 16, 59–66.

Williams, T., 1996. The Iridoviruses. Adv. Virus Res. 46, 345–412.

Xu, X., Zhang, L., Weng, S., Huang, Z., Lu, J., Lan, D., Zhong, X., Yu, X., Xu, A., He, J., 2008. A zebrafish (*Danio rerio*) model of infectious spleen and kidney necrosis virus (ISKNV) infection. Virology 376, 1–12.

Yanong, R.P., Waltzek, T.B., 2010. Megalocytivirus infections in fish, with emphasis on ornamental species. Program in Fisheries and Aquatic Sciences (FA182), University of Florida, 1–7.

Zainathan, S.C., Balaraman, D., Ambalavanan, L., Moorthy, P.K., Palakrishnan, S.K., Ariff, N., 2019. Molecular screening of infectious spleen and kidney necrosis virus in four species of Malaysian farmed ornamental fish. Malays. Appl. Biol. 48, 131–138.

Zainathan, S.C., Johan, C.A.C., Subramaniam, N., Ahmad, A.A., Halim, N.I.A., Norizan, N., 2017. Detection and molecular characterization of Megalocytivirus strain ISKNV in freshwater ornamental fish from Southern Malaysia. AACL Bioflux. 10, 1098–1109.

