## Supplementary figures and images for "Detection and genetic analysis of infectious spleen and kidney necrosis virus (ISKNV) in ornamental fish from non-clinical cases: First report from India"

### Suppl. Fig. 1

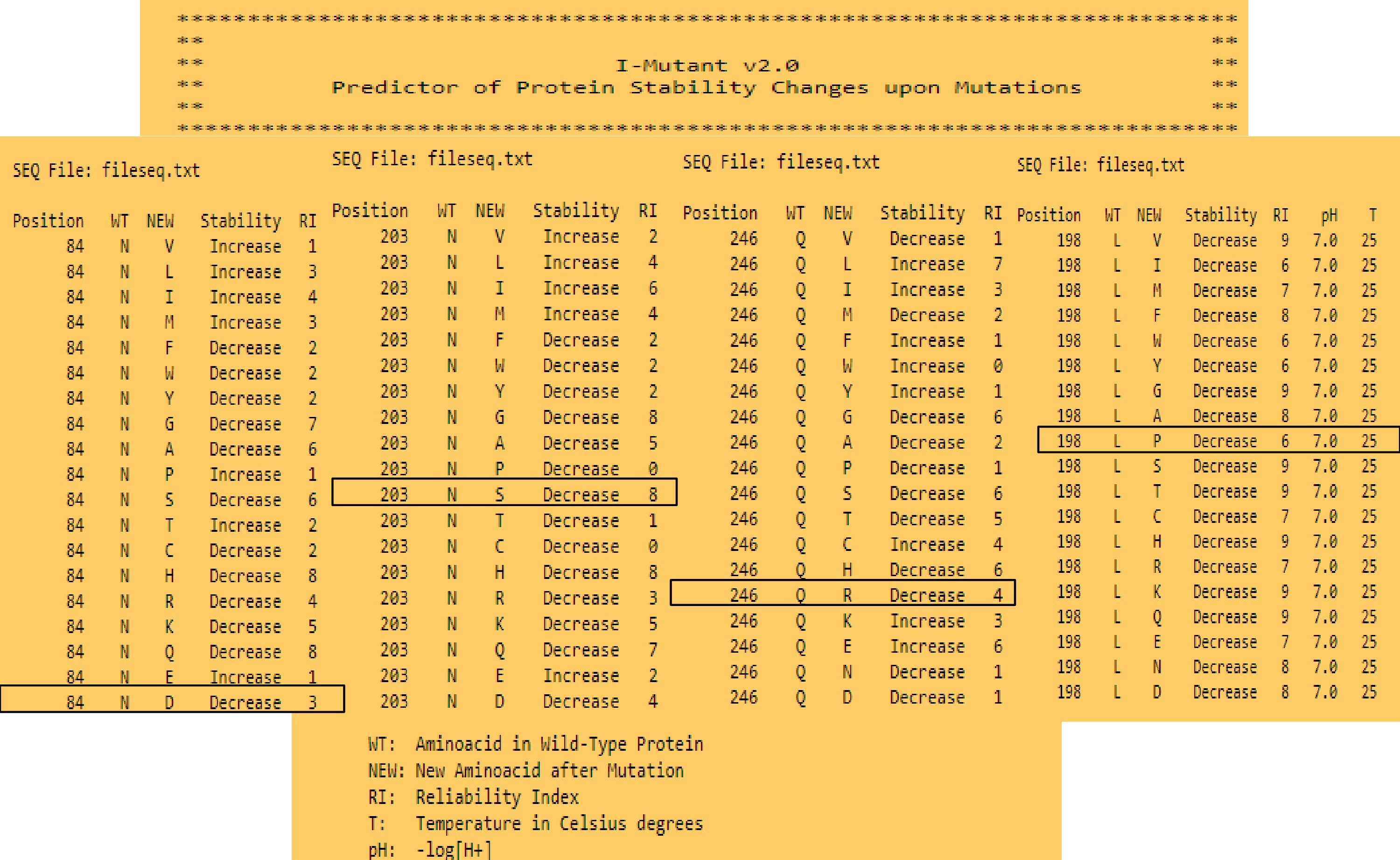

### Suppl. Fig. 2

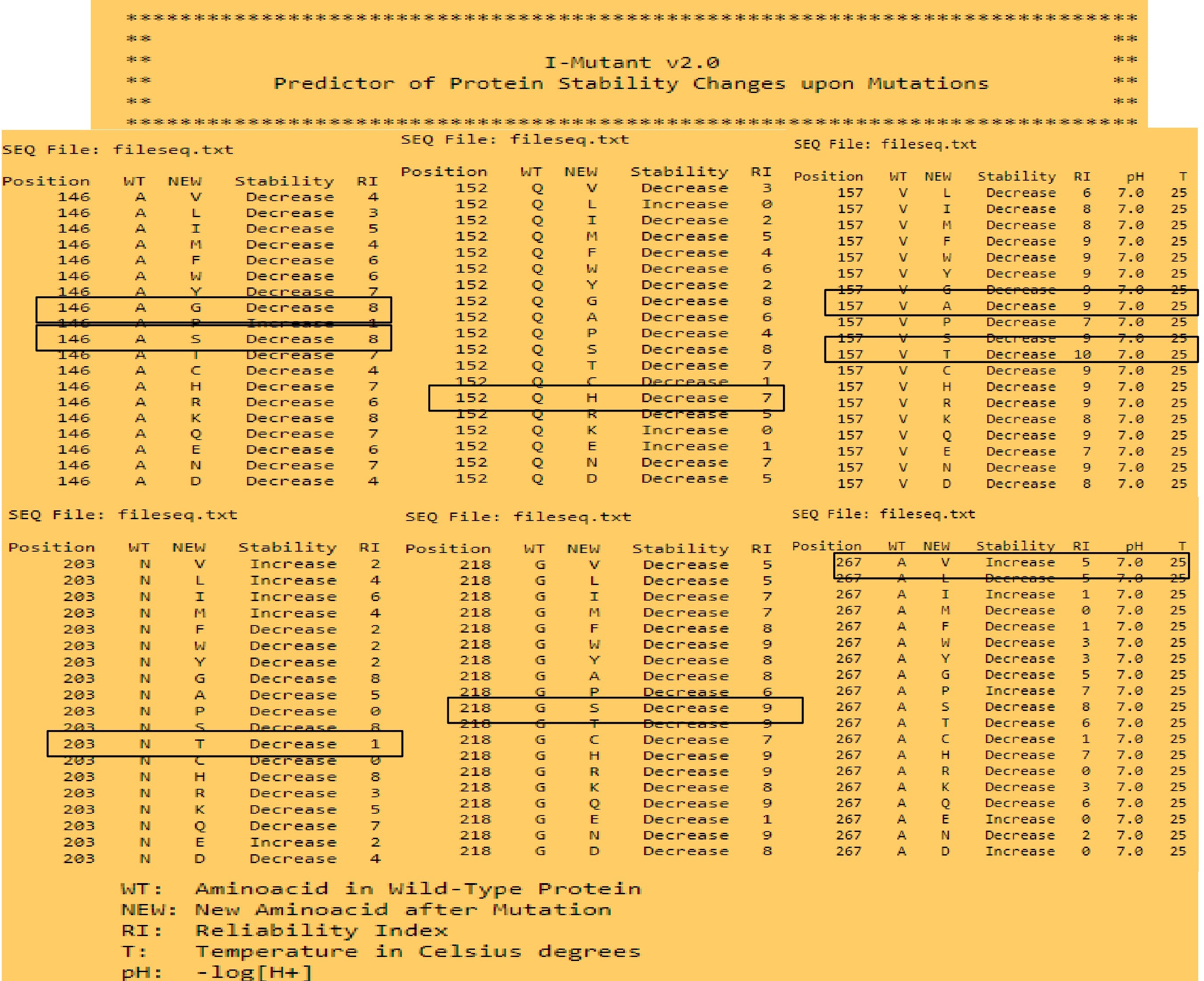
